# Partially dissociable roles of the Orbitofrontal cortex and dorsal Hippocampus in context-dependent (hierarchical) reward prediction and contextual inference in learning

**DOI:** 10.1101/2024.06.19.599779

**Authors:** Sophie Peterson, Jose Chavira, Alex Garcia Arango, David Seamans, Emma Cimino, Ronald Keiflin

**Affiliations:** Department of Psychological & Brain Sciences, University of California, Santa Barbara, CA 93106, USA; Neuroscience Research Institute, University of California, Santa Barbara, CA 93106, USA

**Author notes:** Correspondence should be addressed to R.K.

## Abstract

Reward cues are often ambiguous; what is good in one context is not necessarily good in another context. To solve this ambiguity, animals form hierarchical associations in which the context acts as a gatekeeper in the retrieval of the appropriate cue-evoked memory, ensuring context-appropriate behavior. These hierarchical associative structures also influence future learning by promoting the formation of new context-dependent associations (leading to the inference of context-dependency for new associations). The orbitofrontal cortex (OFC) and the dorsal hippocampus (DH) are both proposed to encode a “cognitive map” that includes the representation of hierarchical, context-dependent, associations. However the causal role of the OFC and DH in the different functional properties of hierarchical associations remains controversial. Here we used chemogenetic inactivations, in rats, to examine the role of OFC and DH in 1) the contextual regulation of performance, and 2) the contextual learning bias conferred by hierarchical associations. We show that OFC is required for both manifestations of hierarchical associations. In contrast, DH contribution appears limited to the contextual learning bias. This study provides novel insight into the different functional properties of context-dependent hierarchical associations, and establishes the OFC as a critical orchestrator of these different contextual effects.

## INTRODUCTION

Stimuli paired with reward are powerful motivational triggers and can elicit anticipatory reward-seeking behaviors. But in a complex world in which the meaning of stimuli is often ambiguous and context-dependent, indiscriminate responding to all reward-signaling cues is not adaptive. Instead adaptive behavior requires the integration of reward cues with the larger context in which these cues are encountered (e.g. the word “pain” means different things in France and the US). In the associative framework, this integration takes the form of hierarchical associations in which the context acts as a gatekeeper in the retrieval of the appropriate cue→outcome association, thereby “setting the occasion” for context-appropriate responses to ambiguous cues^1,2^.

Importantly, the knowledge of contextual rules for stimulus disambiguation (which we refer to as “contextual knowledge”) influences not only the retrieval of established associations (i.e. performance) but also the formation of new associations. Indeed, animals equipped with contextual knowledge are prone to form context-dependent associations during new learning episodes (contextual learning bias) which results in the (inductive) inference of context-dependency for these newly formed associations^3–6.^

This influence of contextual knowledge on performance and learning is a defining feature of cognitive control and is impaired in several neuropsychiatric disorders,including schizophrenia, autism spectrum disorders, post-traumatic stress disorders, and impulse control disorders^7–10^. Yet, despite the relevance of these processes, the neural circuits mediating these manifestations of contextual knowledge remain poorly characterized.

The orbitofrontal cortex (OFC) and the dorsal hippocampus (DH) are both proposed to encode a “cognitive map”, or mental representation of the relationship between events —including the representation of hierarchical, context-dependent, associations^11,12^. This view is largely supported by human fMRI and rodent single-cell recording studies showing encoding of contextual and/or relational information in the OFC and the DH^13–17^. However, the causal role of these structures in the manifestation of contextual knowledge remains uncertain as very few studies examined the neural basis of context-informed hierarchical associations in animal models.

Recent studies in rats showed that inactivation of the OFC, but not the DH, disrupts occasion setting, i.e. the ability of one contextual cue (the occasion setter) to modulate conditioned responding to another (target) cue^18–20^. However, most of these prior investigations focused on unidirectional occasion setting in which the contextual cue either increases or decreases the probability of reinforcement. These preparations leave room for direct associations (positive or negative) between the contextual cue and the outcome. These direct associations might contribute to the ability of the occasion setter to modify responding, casting doubt on the hierarchical nature of associative predictions in these tasks. Moreover, all these prior investigations focused exclusively on one manifestation of contextual knowledge: the retrieval of established associations. The contribution of the OFC and DH to the inference of context-dependency during new learning remains to be determined.

Therefore, the purpose of this study was to examine the potential involvement of the OFC and DH in two manifestations of contextual knowledge: 1) the retrieval of established context-dependent associations and 2) the inference of context-dependency during new associative learning. Rats were trained in a context-dependent Pavlovian discrimination task in which the context exerts opposite influences (positive or negative modulation) on two target cues (i.e. a bidirectional occasion setting task). Following acquisition, we used chemogenetics to silence OFC or DH neurons at two time points: 1) during performance of this learned context-dependent discrimination, and 2) during new associative learning (when animals equipped with contextual knowledge normally infer context-dependency for a newly formed cue-outcome association). Consistent with prior studies, we show that OFC -but not DH-inactivation impairs discrimination performance in a bidirectional occasion setting task. In contrast, both OFC and DH were essential for knowledge-biased acquisition of new context-dependent associations (i.e. inference of context-dependency in new learning).

## RESULTS

Male and female rats were initially trained in the context-dependent discrimination (n=48; 26M + 22F) or the simple discrimination task (n=33; 18M + 15F). In the context-dependent discrimination task, a background visual stimulus (context A) informs the validity of two ambiguously predictive auditory cues, so that one cue (X) is rewarded with sucrose only in presence of A, while the other cue (Y) is rewarded only in the absence of A (A:X+ / X- / A:Y- / Y+). In the simple discrimination task, only one cue (X) is rewarded, independently of the context (A:X+ / X+ / A:Y- / Y-). The two tasks promote the emergence of different associative architectures for reward prediction (**Fig. 1A-C**). Following acquisition of discriminated Pavlovian approach (80 sessions for context-dependent discrimination; 20 sessions for simple discrimination, **Fig. 1D-E**), rats were divided in subgroups of matched performance and injected with AAVs for the expression of the inhibitory DREADD hM4Di, or the control protein mCherry, in the ventral lateral orbitofrontal cortex (OFC) or the dorsal hippocampus (DH) (n = 8-12 / group; 38-50% of females / group), Fig. 1F-G. DREADD-mediated inhibition of the OFC or DH was achieved via systemic injection of CNO (5mg/mg, i.p., 30 min before session).

**Fig. 1.**
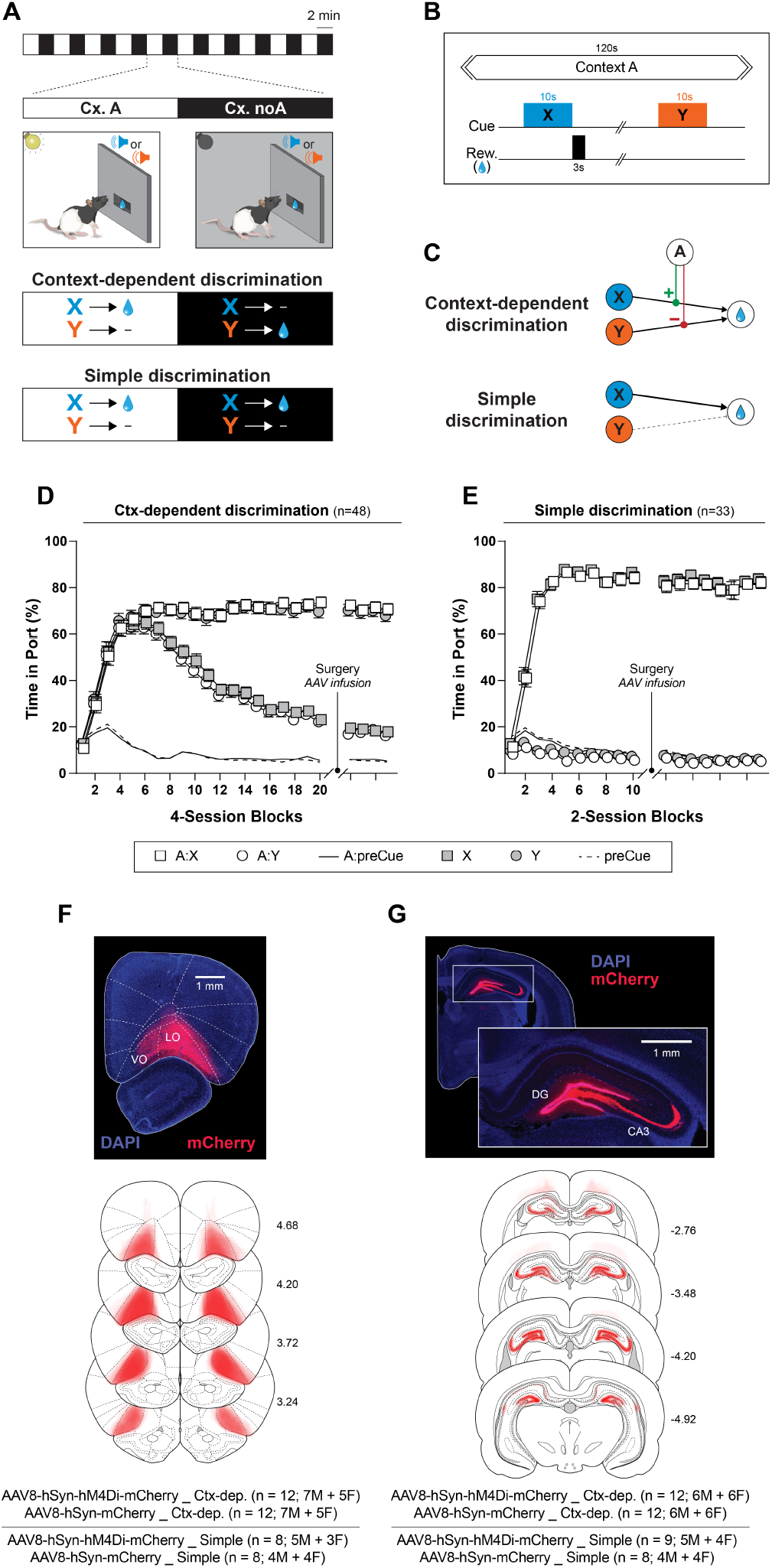
Behavioral tasks and histo-logical verification of virus expression. A-B. Context-dependent and Simple discrimination tasks. During conditioning sessions, the visual background alternates between context A (houselight on; 2 min) and context noA (houselight off; 2 min). Embedded in these contexts cycles, brief auditory cues (X and Y; 10s each) are presented and potentially followed by a liquid sucrose reward (B). In the context-dependent discrimination task, the predictive status of these cues is informed by the background visual context (X is rewarded only in presence of A whereas Y is rewarded only in absence of A). In the simple discrimination task, only once cue (X) is rewarded and the background context is irrelevant.C. Associative structures engaged by the two tasks. Context-dependent discrimination relies on hierarchical associations in which the context acts as a “gatekeeper” in the retrieval of the appropriate cued prediction.D-E. Pavlovian approach behavior (time in port) evoked by the different trial types (A:X / X / A:Y / Y) across the acquisition period, for rats trained in the context-dependent (D) or simple discrimination (E) task. Dashed line show time in port in the 10s prior to cue presentation.F-G. Post-mortem verification of virus expression in the orbitofrontal cortex (F) or the dorsal hippocampus (G). Top: representative fluorescent image of hM4Di-mCherry expression. Bottom reconstructed representation of viral expression (mCherry and hM4Di-mCherry subjects combined). Note that virus expression in the DH was largely restricted to the dentate gyrus (DG) and CA3, with very sparse expression in CA1.

### OFC inactivation disrupts the contextual regulation of cued reward seeking

We first examined the effect of DREADD-mediated OFC inactivation on context-dependent or simple discrimination performance (**Fig. 2A**). For each OFC virus and training condition, cue-evoked responding (i.e. time in port) following vehicle or CNO was analyzed with 3-way RM-ANOVAs (cue x context x injection). As expected, rats trained in the Ctx-dep. discrimination task displayed a context-sensitive response to the reward cues (Cue x Context interaction: hM4Di: F(1,11)= 99.32, P < 0.001; mCherry: F(1,11) = 181.130, P < 0.001). In contrast, animals trained in the simple discrimination task responded selectively to the reward cue (Cue main effect: hM4Di: F(1,7) = 164.328, P < 0.001, mCherry: F(1,7) = 130.490, P < 0.001) and this response was independent of the context (Cue x Context interaction: hM4Di: F (1,7) = 0.082, P = 0.783; mCherry: F(1,7) = 0.951, P = 0.362). Critically, only animals trained in the Ctx dep. discrimination task and expressing the inhibitory hM4Di DREADD showed disrupted performance following CNO injection (Cue x Context x Injection interaction: F(1,11) = 58.439, P < 0.001). DREADD-mediated OFC inactivation selectively increased responding to nonrewarded (contextually irrelevant) cues (Ps ≥ 0.001; post-hoc t-tests) while responding to rewarded cues remained high and not affected by OFC inactivation (Ps ≥ 0.104). Critically, CNO injection had no effect in rats lacking the inhibitory DREADD (mCherry controls) or in rats trained in the simple discrimination task, regardless of virus expression (no main effect, or interaction with Injection: Ps ≥ 0.061; **Fig. 2B**).

**Fig. 2.**
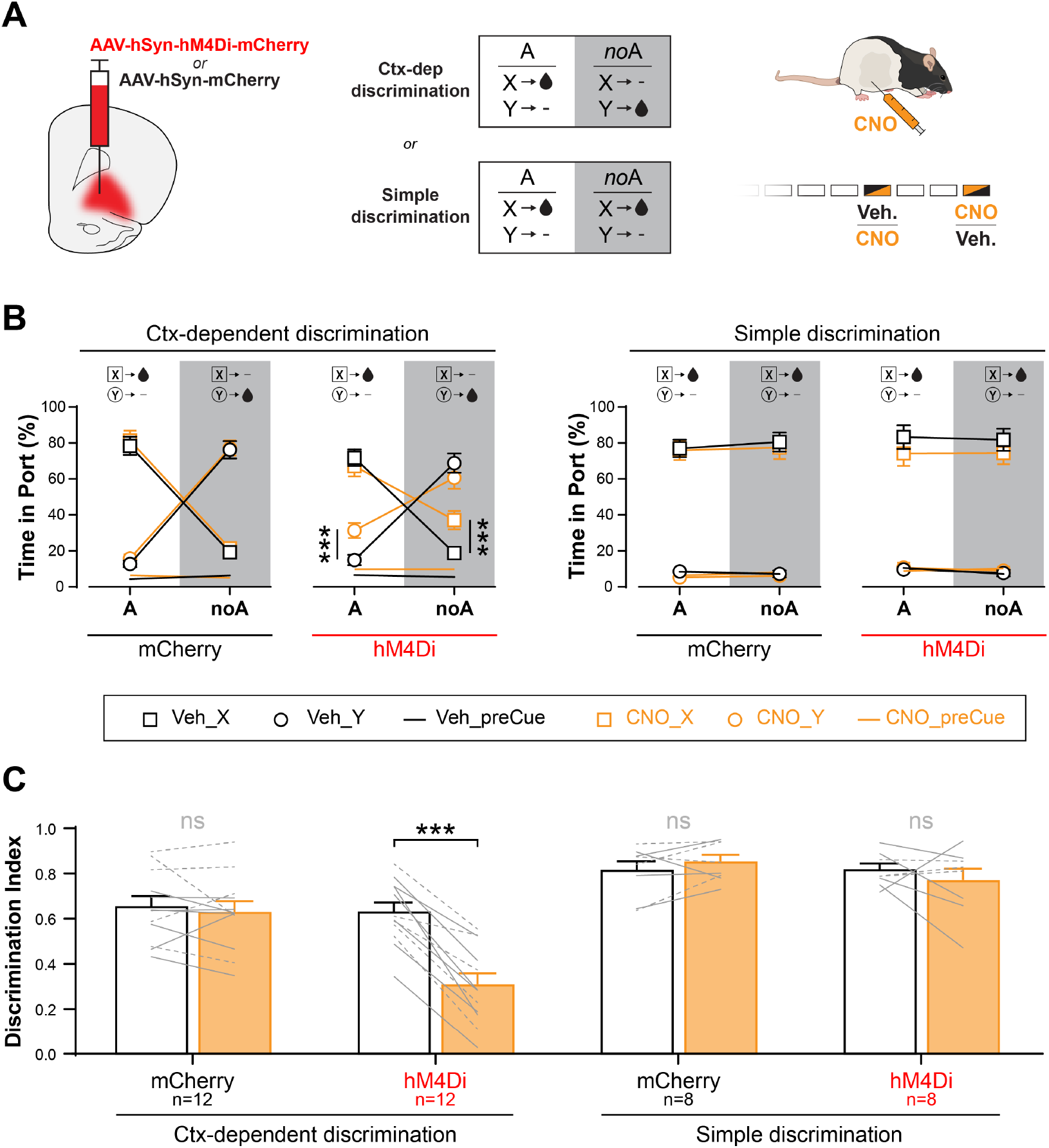
OFC inactivation selectively impairs the contextual regulation of cued reward seeking. **A.** Experimental design: rats expressing the inhibitory DREADD hM4Di or the control protein mCherry in the OFC were trained in the context-dependent, or the simple discrimination task. The effect of acute CNO (5mg/ml, ip) or saline (vehicle) on discrimination performance was evaluated for all rats, in two separate sessions (order of injection counterbalanced). **B**. Pavlovian approach behavior (time in port) evoked by the different trial types (A:X / X / A:Y / Y during saline and CNO sessions. For rats trained in the context-dependent discrimination task, the hM4Di/CNO-mediated silencing of the OFC increased contextually inappropriate responding to non-rewarded cues. **C**. Discrimination index during Vehicle and CNO sessions. CNO injection reduced discrimination index (i.e. reduced discrimination accuracy) only in rats expressing hM4Di in the OFC and trained in the context-dependent task. Solid gray lines = males; dashed gray lines = females. *** P < 0.001 Post-hoc paired t-tests (Veh. vs CNO). Error bars = s.e.m.

To facilitate direct comparison between groups, we calculated for each animal a discrimination index, defined as the difference between time-in-port during rewarded and nonrewarded trials divided by the sum (DI= 0: no discrimination; DI = 1: perfect discrimination). A 3-way mixed-design ANOVA (Task x Virus x Injection) conducted on this discrimination index confirmed a task- and virus-sensitive effect of CNO (Task x Virus x Injection interaction: F(1,36) = 7.352, P = 0.010) as CNO decreased discrimination accuracy only in rats trained in the Ctx-dependent task and expressing hM4Di (P ≥ 0.001); CNO had no effect in any other group (Ps ≥ 0.266; **Fig 2C**).

### DH inactivation spares the contextual regulation of cued reward seeking

Next, we examined the effect of DREADD-mediated DH inactivation on context-dependent or simple discrimination performance (**Fig. 3A**). Rats trained in the Ctx-dep. discrimination task and expressing hM4Di in the DH showed a modest but significant increase in responding (time in port) after CNO-induced DH inactivation (main Injection effect: F(1,11) = 10.297, p = 0.008). This increase in responding was more pronounced in the CxA (Context x Injection: F(1,11) = 4.924, p = 0.048) but affected rewarded and nonrewarded cues equally (no Cue x Injection, or Cue x Context x Injection interaction, Ps ≥ 0.325; **Fig. 3B**). Importantly, this small generalized increase in time-in-port did not translate into a decrease in discrimination accuracy as measured by the discrimination index (T(11) = 1.137; P = 0.280; targeted paired t-test). CNO injection had otherwise no effect on time in port for any other group (no main effect or interaction with Injection: Ps ≥ 0.335), and overall, CNO had no effect on discrimination index for any DH virus groups (no main effect or interaction with Injection: Ps ≥ 0.386; **Fig. 3C**).

**Fig. 3.**
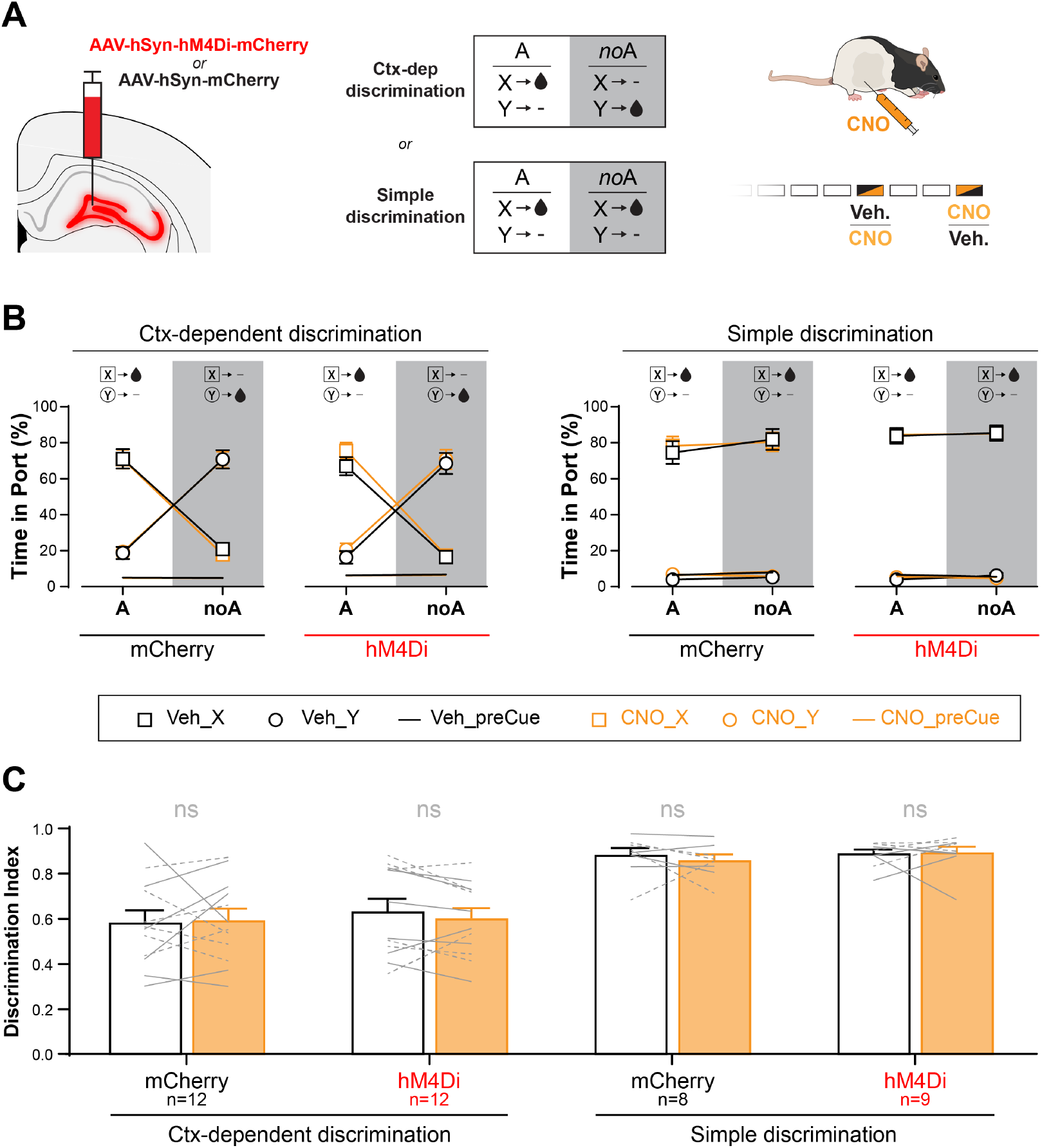
DH inactivation spares the contextual regulation of cued reward seeking. **A.** Experimental design: rats expressing the inhibitory DREADD hM4Di or the control protein mCherry in the OFC were trained in the context-dependent, or the simple discrimination task. The effect of acute CNO (5mg/ml, ip) or saline (vehicle) on discrimination performance was evaluated for all rats, in two separate sessions (order of injection counterbalanced). **B**. Pavlovian approach behavior (time in port) evoked by the different trial types (A:X / X / A:Y / Y during vehicle and CNO sessions. **C**. Discrimination index during Vehicle and CNO sessions. CNO injection did not reduce discrimination accuracy in any experimental group. Solid gray lines = males; dashed gray lines = females. Error bars = s.e.m.

### Prior knowledge of contextual rules for reward prediction promotes the inference of context-dependency in new learning

Prior studies showed that animals and humans previously trained in context-dependent discrimination tasks were prone to infer a context-dependent rule during new learning episodes^3–5^. This phenomenon has been captured by several computational or neural network models^3,21,22^ (**Fig. S1**). To determine if this phenomenon could be replicated in our preparation, surgery-naïve rats previously trained in the context-dependent discrimination task were introduced to a new cue Z during 3 training sessions. Critically, this new cue was presented only in presence of the contextual feature A, and always rewarded. These A:Z+ trials (10 per session) were interspersed with regular trials featuring the original cues and their usual consequences (A:X+ / X- / A:Y- / Y+). The proportion of these trials was arranged to maintain an equal number of rewards in both contexts. These sessions introducing the cue Z were themselves interleaved with regular training sessions featuring only the original cues. This protocol ensured that new associative learning occurred against the backdrop of stable prior knowledge. Finally, we assessed to which extent responding to this new cue Z was modulated by the context in a final non-rewarded probe test. Although context-dependency was never explicitly confirmed for this new cue, rats previously trained in the context-dependent discrimination task generalized this contextual rule. This resulted in context-sensitive responding to this new cue Z, as revealed in the final probe test (A:Z vs. Z: T(11) = 5.406; P<0.001; Fig. S1). This inference of context-dependency was not observed in task-naïve rats or in rats previously trained in the simple discrimination task and exposed to similar new cue training. Context-sensitive responding to cue Z was also not observed if that cue was not rewarded during the previous sessions (i.e. no acquired equivalence between Z and Y).

### OFC inactivation prevents the inference of context-dependency in new learning

Having shown that prior knowledge of contextual rules biases new learning and promotes the formation of context-dependent associations, we next examined the contribution of the OFC in this process (**Fig. 4**). Rats previously trained in the context-dependent discrimination task and expressing hM4Di or mCherry in the OFC (same rats as in prior OFC experiment) were introduced to a new reward cue Z. As previously described, the new cue Z was always accompanied by the contextual feature A and always rewarded. All rats received injections of the DREADD agonist CNO prior to the sessions introducing the new cue Z (effectively inactivating OFC in hM4Di-expressing rats during this new associative learning episode; **Fig.4A**). On the first presentation of cue Z, anticipatory reward seeking was virtually absent; this indicates that this new cue was easily distinguishable from the original cues (X and Y) and there was no spontaneous generalization between cues. Responding to cue Z then quickly emerged in both groups and there was no effect of OFC inactivation on the acquisition of this new conditioned response (Session: F(2, 42) = 27.999; P < 0.001; no main effect or interaction with Virus Ps ≥ 0.103; **Fig. 4B**). Note that the introduction of this new cue Z produced a small and transient reduction in the discrimination accuracy for the original cues X and Y (F(1, 21) = 24.663, P < 0.001), suggesting minor interference between prior knowledge and new learning (**Fig. 4C**). Surprisingly, OFC inactivation — which earlier disrupted the contextual regulation of behavior— had no effect on response accuracy in these sessions (no main effect or interaction with Virus, Ps ≥ 0.142). This suggests that the introduction of the new cue, and/or the repeated OFC inactivation, promoted the recruitment of additional brain regions which might have compensated for the absence of functional OFC. Finally, anticipatory reward seeking evoked by all cues was tested in both contexts in two non-rewarded probe tests (test #1: A:Z / Z; test #2: A:X / X / A:Y / Y). As previously shown for surgery-naïve animals, rats trained in the context-dependent task and lacking the inhibitory DREADD (mCherry controls) demonstrated a contextual learning bias and generalized context-dependency to the new cue Z (A:Z > Z). Inactivation of the OFC during the new learning phase abolished this effect (**Fig. 4D**). Indeed, rats expressing hM4Di in the OFC (for which OFC was inactivated during learning) displayed robust responding to cue Z during the probe test (when OFC was again fully functional) but this behavior was not modulated by the context (Context x Virus: F(1, 21) = 5.447, P = 0.030; A:Z vs. Z: mCherry: P= 0.003, hM4Di: P = 0.990). Contextual modulation was not observed in rats trained in the simple discrimination task, regardless of OFC virus expression (no main effect of Context, Virus, or interaction between these factors, Ps ≥ 0.611; **Fig. 4I**).

**Fig. 4.**
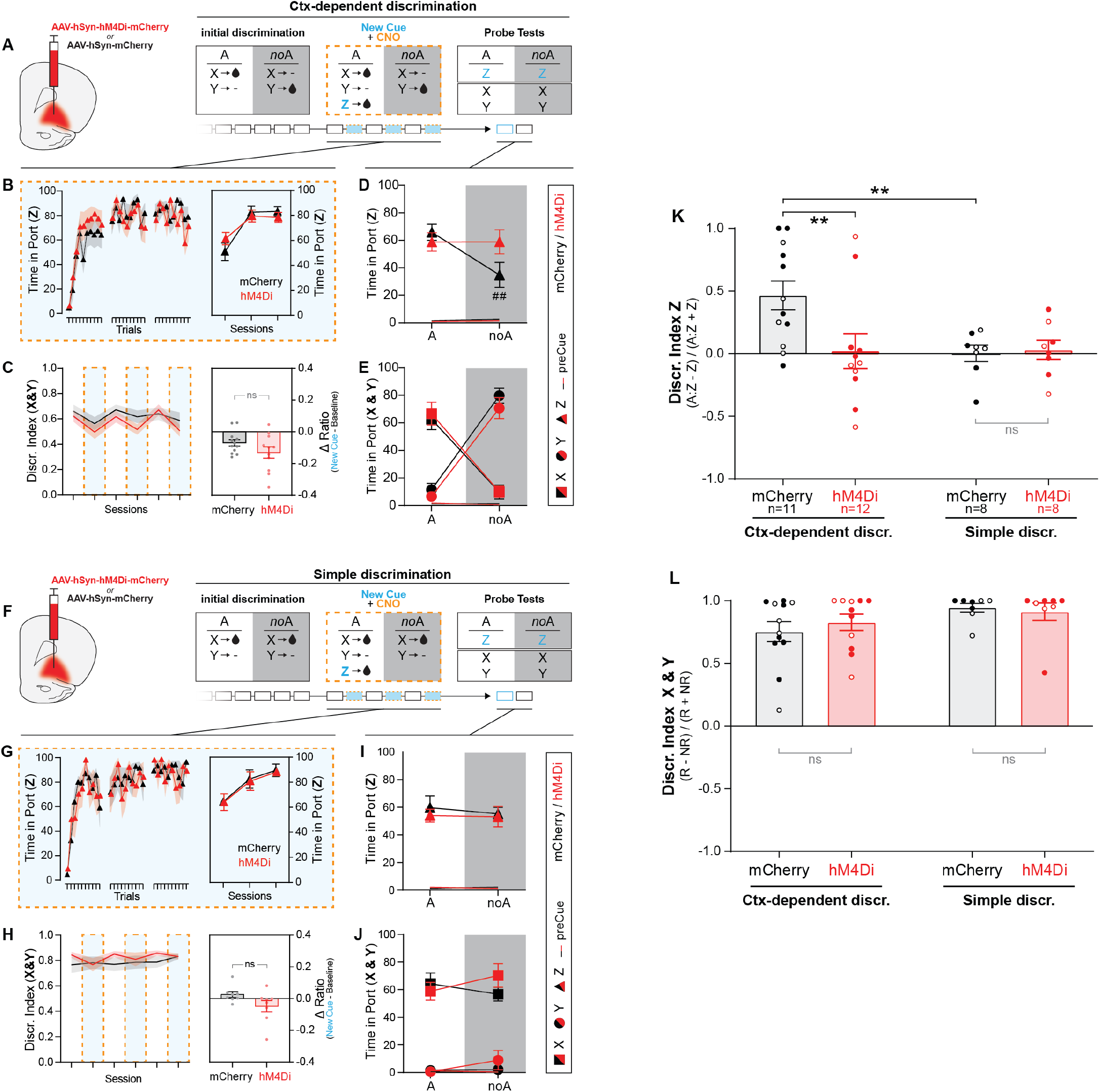
OFC inactivation prevents the inference of context-dependency during new associative learning. **A & F**. Experimental design: rats expressing hM4Di or mCherry in the OFC were trained in the context dependent discrimination (**A**) or the simple discrimination task (**F**). Following initial discrimination training, a new cue (Z) was introduced —always in the presence of the contextual feature A, and always rewarded. Those (A:Z+) trials were interspersed with regular trials featuring the original cues X and Y and their regular programmed outcomes. All rats received CNO injections prior to the sessions introducing the new cue Z. The ability of the context to modulate cue-evoked responding was then assessed in a final, drug-free, probe test during which all cues were presented in both contexts. **B & G**. Acquisition of conditioned approach behavior to the new cue Z, across trials (left) and across sessions (right), for rats trained in the context-dependent (**B**) or simple discrimination task (**G**). **C & H** Discrimination index for the original target cues (X & Y) during the new cue (+CNO) sessions and the alternating regular (drug-free) sessions. The introduction of a the new cue led to a small acute reduction in discrimination accuracy in rats trained in the context-dependent (**C**), but not the simple discrimination task (**H**), regardless of virus expression. Inset: difference score (discr. index during regular session – discr. index during new cue session). **D & I**. Conditioned approach behavior evoked by cue Z during the final probe test. Control (mCherry) rats trained in the context dependent discrimination showed a contextual learning bias and inferred context-dependency for cue Z (i.e. more responding to Z presented in context A vs noA). This effect was abolished in hM4Di rats who learned about Z while their OFC was temporarily inactivated (**D**). No context dependency for cue Z was inferred in the rats trained in the simple discrimination task, regardless of virus expression (**I**). **E & J**. Conditioned approach behavior evoked by the original cues X and Y during the final probe test, for rats trained in the context-dependent (**E**) or simple discrimination task (**J**). **K**. Contextual modulation index for cue Z during the final probe test. **L**. Discrimination index for the original cues X and Y during the final probe test. Filled symbols: males; empty symbols: females. ##: P < 0.01 Post-hoc paired t-tests (AZ vs Z). **: P < 0.01 Post-hoc independent t-tests. Error bars = s.e.m.

To directly compare the contextual modulation of cue Z in all groups, we calculated for each animal a context modulation index ([A:Z – Z] / [A:Z + Z]) (**Fig. 4K**). A 2-way ANOVA (Task x Virus) conducted on this index showed that rats previously trained in the context-dependent task and with functional OFC during new learning (Ctx-dep. mCherry) stood different from all other groups as only these animals displayed context-sensitive responding to the cue Z (Task x Virus: F(1, 35) = 4.201, P = 0.048; Ctx-dep hM4Di vs. all other groups: Ps ≥ 0.007). No effect of virus expression/ OFC inactivation was observed on the discrimination accuracy to the original cues X and Y during the final probe test (**Fig. 4E, J, L**).

### DH inactivation prevents the inference of context-dependency in new learning

Next, we examined the contribution of the DH in the inference of context-dependency during new learning, using the same procedure as previously described. Rats previously trained in the context-dependent discrimination task and expressing hM4Di or mCherry in the DH (same rats as in prior DH experiment) were introduced to a new reward cue Z accompanied by the contextual feature A (**Fig. 5**). Rats with functional DH during this new learning phase (mCherry group) later displayed an inferred context-sensitive response to this cue Z during the final probe test (A:Z > Z). This effect was abolished by DREADD-mediated DH inactivation during the new learning phase (Context x Virus: F(1, 22) = 7.474, P = 0.012; A:Z vs. Z: mCherry: P < 0.001; hM4Di: P = 0.956; **Fig. 5D**). Contextual modulation was not observed in rats trained in the simple discrimination task, regardless of DH virus expression (no main effect of Context, Virus, or interaction between these factors, Ps ≥ 0.101; **Fig. 5I & K**). No effect of virus expression/ DH inactivation was observed on the discrimination accuracy to the original cues X and Y during the final probe test (**Fig. 5E,J, L**). In summary, the DH, while not required for the expression of context-dependent discrimination (as shown above), was essential for the inference of context-dependency during new learning episodes.

**Fig. 5.**
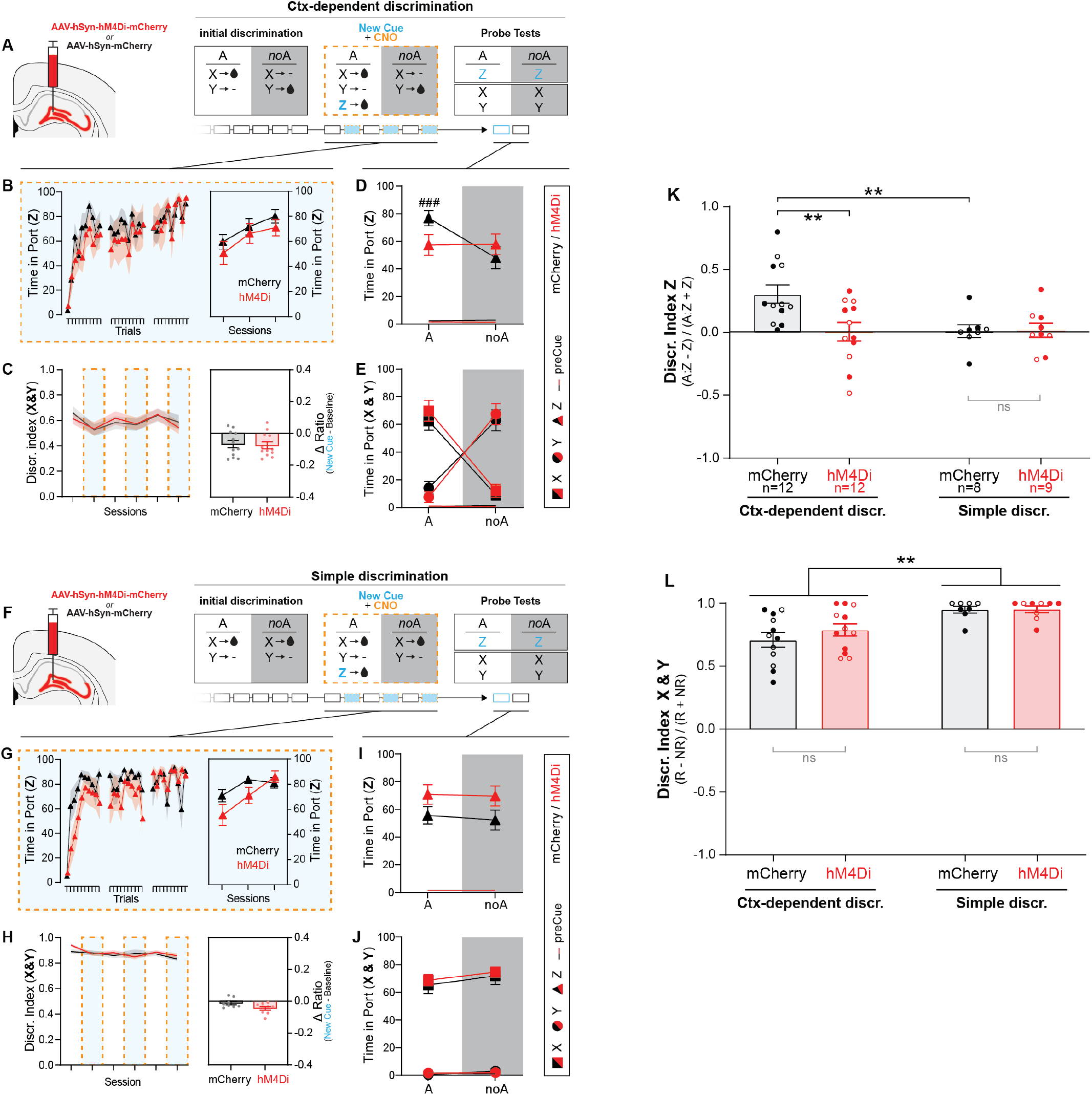
DH inactivation prevents the inference of context-dependency during new associative learning. **A & F**. Experimental design: rats expressing hM4Di or mCherry in the DH were trained in the context dependent discrimination (**A**) or the simple discrimination task (**F**). Following initial discrimination training, a new cue (Z) was introduced —always in the presence of the contextual feature A, and always rewarded. Those (A:Z+) trials were interspersed with regular trials featuring the original cues X and Y and their regular programmed outcomes. All rats received CNO injections prior to the sessions introducing the new cue Z. The ability of the context to modulate cue-evoked responding was then assessed in a final, drug-free, probe test during which all cues were presented in both contexts. **B & G**. Acquisition of conditioned approach behavior to the new cue Z, across trials (left) and across sessions (right), for rats trained in the context-dependent (**B**) or simple discrimination task (**G**). **C & H**. Discrimination index for the original target cues (X & Y) during the new cue (+CNO) sessions and the alternating regular (drug-free) sessions. The introduction of a the new cue led to a small acute reduction in discrimination accuracy in rats trained in the context-dependent (**C**), but not the simple discrimination task (**H**), regardless of virus expression. Inset: difference score (discr. index during regular session – discr. index during new cue session). **D & I**. Conditioned approach behavior evoked by cue Z during the final probe test. Control (mCherry) rats trained in the context dependent discrimination showed a contextual learning bias and inferred context-dependency for cue Z (i.e. more responding to Z presented in context A vs noA). This effect was abolished in hM4Di rats who learned about Z while their DH was temporarily inactivated (**D**). No context dependency for cue Z was inferred in the rats trained in the simple discrimination task, regardless of virus expression (**I**). **E & J**. Conditioned approach behavior evoked by the original cues X and Y during the final probe test, for rats trained in the context-dependent (**E**) or simple discrimination task (**J**). **K**. Contextual modulation index for cue Z during the final probe test. **L**. Discrimination index for the original cues X and Y during the final probe test. Filled symbols: males; empty symbols: females. ###: P < 0.001 Post-hoc paired t-tests (AZ vs Z). **: P < 0.01 Post-hoc independent t-tests. Error bars = s.e.m.

### OFC or DH inactivation does not protect context-dependent predictions against retroactive interference

Our results thus far indicate that OFC or DH inactivation does not affect the acquisition of simple (direct) cue-reward associations, but prevents the integration of this new learning with the prior knowledge of contextual rules. Next, we examined the boundaries of this effect by assessing the effect of OFC or DH inactivation on retroactive interference _another form of interaction between prior knowledge and new learning. Theoretical models predict that repeated rewarded presentation of cue Z in a context-independent manner (A:Z+ / Z+) should interfere with (and disrupt) prior knowledge of context-dependent associations, but not direct context-independent associations (**Fig. S2**). This was confirmed in surgery-naïve rats; repeated reinforcement of cue Z in both contexts (no other cues presented during those sessions) diminished discrimination accuracy for the original target cues in rats originally trained in the context-dependent, but not the simple discrimination task, as revealed in a final probe test (**Fig. S2**). We then used a similar behavioral procedure (using the same rats as in prior experiments) to determine how DREADD-mediated OFC or DH inactivation during the A:Z+ / Z+ sessions affects retroactive interference (OFC: **Fig. 6;** DH: **Fig. 7**). As previously shown in surgery-naïve rats, OFC or DH AAV-injected rats trained in the context-dependent discrimination task —but not the simple discrimination task— saw a decrease in discrimination accuracy post A:Z+/Z+ sessions. This effect was not affected by OFC or DH inactivation. Together, these results indicate that while functional OFC and DH are essential for the interaction between new learning and prior contextual knowledge; silencing, or putting these structures offline during learning does not isolate existing knowledge from disrupting new information.

**Fig. 6.**
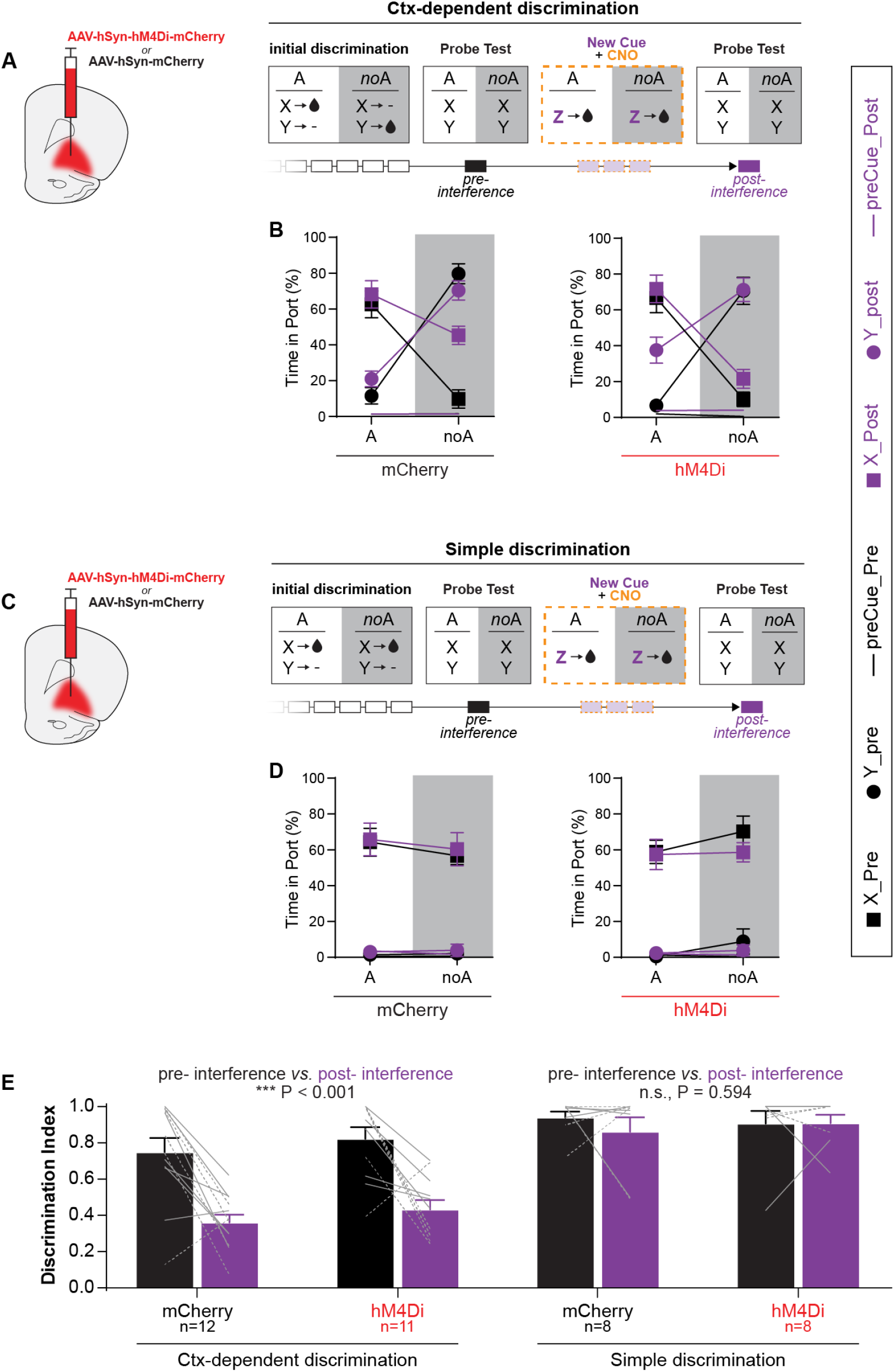
OFC inactivation does not protect context-dependent associations from interfering information. **A & C**. Experimental design. Rats expressing hM4Di or mCherry in the OFC were trained in the context dependent discrimination (**A**) or the simple discrimination task (**C**). Following initial discrimination training, a new reward cue (Z) was introduced for 3 sessions. This new cue was always rewarded regardless of context (A:Z+ / Z+), contradicting the principle of context-dependency for rats trained in the context-dependent discrimination task. All rats received CNO injections prior to these sessions. The effect of this new information on initial associative memories was evaluated in two probe tests (pre- and post-interference by Z). **B & D**. Pavlovian approach behavior (time in port) during presentations of the original cues X and Y, during the probe tests. Rats trained in the context dependent discrimination task showed a reduction of discrimination accuracy following the A:Z+ / Z+ sessions, regardless of virus expression (**B**). Rats trained in the simple discrimination task maintained high discrimination accuracy following A:Z+ / Z+ sessions (**D**). **E**. Discrimination index during the probe tests. Solid gray lines = males; dashed gray lines = females. Error bars = s.e.m.

**Fig. 7.**
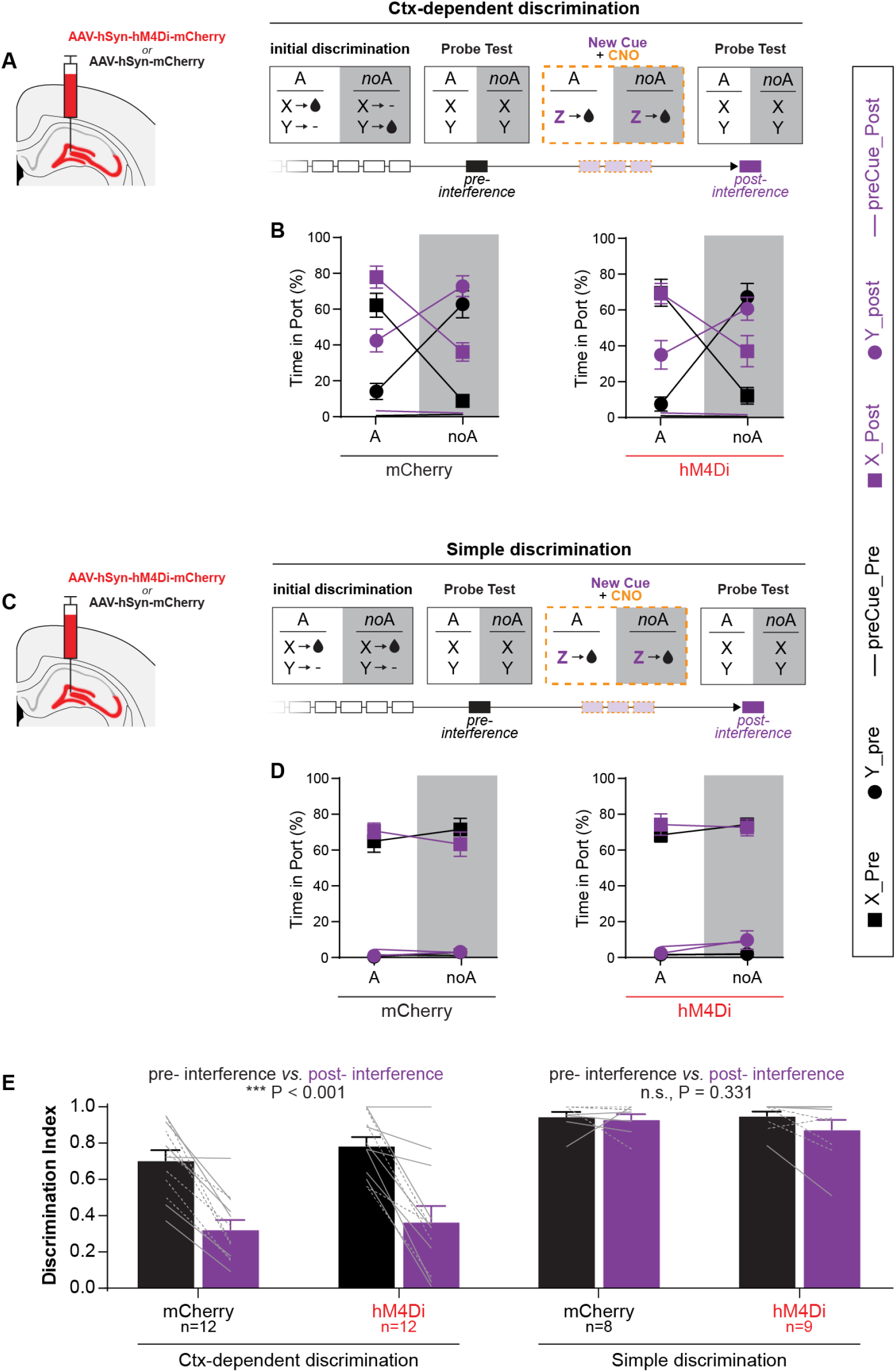
DH inactivation does not protect context-dependent associations from interfering information. **A & C**. Experimental design. Rats expressing hM4Di or mCherry in the DH were trained in the context dependent discrimination (**A**) or the simple discrimination task (**C**). Following initial discrimination training, a new reward cue (Z) was introduced for 3 sessions. This new cue was always rewarded regardless of context (A:Z+ / Z+), contradicting the principle of context-dependency for rats trained in the context-dependent discrimination task. All rats received CNO injections prior to these sessions. The effect of this new information on initial associative memories was evaluated in two probe tests (pre- and post-interference by Z). **B & D**. Pavlovian approach behavior (time in port) during presentations of the original cues X and Y, during the probe tests. Rats trained in the context dependent discrimination task showed a reduction of discrimination accuracy following the A:Z+ / Z+ sessions, regardless of virus expression (**B**). Rats trained in the simple discrimination task maintained high discrimination accuracy following A:Z+ / Z+ sessions (**D**). **E**. Discrimination index during the probe tests. Solid gray lines = males; dashed gray lines = females. Error bars = s.e.m.

## DISCUSSION

Context-dependent associative structures, or context-informed cognitive maps, are thought to reflect the hierarchical nature of predictive representations^1,23^. These context-informed hierarchical knowledge structures (here referred to as “contextual knowledge”) support flexible behavior in at least two ways: 1) by gating the retrieval of associative memories, context resolves the meaning of ambiguous cues and promotes context-appropriate responses to these cues (contextual regulation of performance) and 2) contextual knowledge biases learning and promotes the inference of generalized context-dependency for newly formed associations (contextual learning bias). Here we investigated the role of the OFC and DH is these two manifestations of contextual knowledge. Consistent with OFC’s proposed role in encoding context-dependent predictive relationships, we found that OFC was critical to both manifestations of contextual knowledge. Indeed, chemogenetic inactivation of the OFC disrupted the contextual regulation of performance and abolished the contextual learning bias normally displayed by rats previously trained in the context-dependent discrimination task. In contrast, DH’s contribution to contextual knowledge appears restricted to the inference of context-dependency during new learning (contextual learning bias) with seemingly no implication of the DH in the contextual regulation of performance.

This work extends recent studies implicating the OFC in unidirectional occasion setting^18,19^. In those preparations, one stimulus –the occasion setter (OS) – increases (or decreases) the probability of reinforcement of another conditioned stimulus (CS), therefore increasing (or decreasing) the magnitude of cue-evoked responses. This influence of occasion setters on cue-evoked responses may reflect the hierarchical gating of associative memories, but may also reflect direct associations (positive or negative) between the occasion setter and the reinforcing outcome^24,25^. Therefore, although OFC inactivations were found to disrupt occasion setting, the possibility of a summation between OS- and CS-evoked predictions in those earlier studies cast doubts on OFC’s role in the hierarchical (contextual) control of reward predictions^26,27^. In this study, for rats trained in the context-dependent discrimination task, the context (A) can be conceived as a bidirectional occasion setter, simultaneously increasing and decreasing the predictive validity of two distinct target cues (A:X+ / X-/ A:Y- / Y+). Critically, associative summation cannot account for context-dependent reward seeking in this task. Therefore, our study provides definitive proof of OFC’s involvement in the hierarchical (contextual) control of reward predictions.

The DH, despite its well established role in context processing (particularly spatial and/or multimodal “global” contexts), is not essential for all forms of contextual behaviors^28–30^. Consistent with prior reports, we show here (using a unimodal “local” context) that reversible DH inactivation does not impair behavioral performance in a context-dependent discrimination task^19,31^. This negative result should however be interpreted with caution. Although it clearly demonstrates that contextual control over conditioned appetitive responses is possible without functional DH (after chemogenetic inactivation), it does not rule out a participation of the DH in contextual control earlier in training, before system consolidation^32^, and/or under normal physiological conditions. Indeed, the slow time course of pharmacological chemogenetic inactivations allows for the compensatory recruitment of extra-hippocampal circuits, possibly hiding DH’s normal involvement in the task^33,34^ .

Consistent with several human and rodent studies, we show here that previously acquired knowledge profoundly influences the content and neural substrates of subsequent learning^5,35–37^. It took∼4 months of training (∼80 sessions) for rats to establish stable context-dependent discrimination, yet once trained, rats spontaneously and rapidly inferred context-dependency for a new cue-reward association, even in absence of explicit information to confirm this belief. This influence of prior knowledge on learning might be interpreted as a form of schema assimilation. In this view, the knowledge of a context-dependent rule for cued reward predictions constitutes a memory schema, i.e. a knowledge structure acting as a scaffold for the assimilation of new information. Previous studies highlighted the importance of spatial memory schemas in the acquisition of new item-place associations in rodents, and demonstrated the critical role of the hippocampus and neocortex in this schema assimilation process^36^. Here we highlight the importance of associative (relational) schemas in new cue-reward learning. Moreover we show that the contribution of the DH to schema assimilation is not limited to spatial schemas, but extend to associative schemas more generally. Overall, our findings that OFC or DH inactivation abolish the induction of context-dependent learning are consistent with the proposal that hippocampal-neocortical interactions mediate schema assimilation^38^. The complementary learning systems theory proposes that the hippocampus rapidly encodes context-bounded episodic information that is then gradually integrated into existing knowledge structures in the neocortex during the system consolidation process^39^. Inactivation of the DH or the OFC would then prevent the initial episodic encoding of the new cue-reward pairing, or the post-session integration of this episodic information, respectively. Alternatively, or in addition, the OFC might not only receive information from the hippocampus, but might also modulate hippocampal learning processes by gating the flow of information to the hippocampus (via parahippocampal projections)^16^. This might allow the OFC to direct attention to the features of the environment that are deemed most relevant based on prior experiences and existing knowledge. Further studies are required to determine the mechanisms of this DH-OFC interaction (what is the direction of this interaction, what pathways are involved, and when in the acquisition or consolidation process are these pathways involved?). Note that direct connections between the OFC and DH are extremely sparse/nonexistent, therefore DH-OFC interactions likely involve a complex, multisynaptic circuit^11,40–45^. Questions also remain regarding the degree of abstraction of the cognitive map acquired by rats in our tasks (does prior contextual knowledge promote context-dependent learning only within the training context and for cues/outcomes of the same modality/valence as in training, or is context-dependent learning promoted more generally?)

Importantly, our findings can help reconcile apparent discrepancies between rodent and human associative learning strategies. Whereas rodents typically approach feature positive (A:X+ / X-) or negative (A:X- / X+) discrimination problems using simple, non-hierarchical associative strategies (resorting to hierarchical strategies only when simpler strategies fail to establish accurate predictions), humans spontaneously approach these problems using hierarchical associative strategies^46,47^. Our findings suggest that differences in pre-existing schema, rather than fundamental species differences in learning processes might account for this discrepancy (task-naïve rodent lacking relevant schema, unlike human participants who are well-aware of the possibility of hierarchical relationships).

We found that the OFC contributes to both the contextual regulation of performance and the contextual learning bias. However, these two manifestations of contextual knowledge appear to be partially dissociable. Although OFC inactivation reduced both context-dependent performance and contextual learning bias, the magnitude of these effects were not correlated (**Fig. S3**). Moreover, OFC inactivation abolished contextual learning bias at a time when this inactivation no longer had a detectable effect on performance. The reason for the seemingly temporary role of the OFC on the contextual regulation of performance remains to be determined. One hypothesis is that the introduction of the novel cue Z, and/or to the repeated OFC inactivation promoted the recruitment of additional brain regions which might have compensated for the absence of functional OFC. Overall, our results strongly suggests that the OFC mediates the effects of context on performance and learning via different circuits. For instance, orbitofrontal top-down projections to the striatum might mediate the contextual regulation of performance (top-down regulation of cue’s motivational effects) while indirect, multisynaptic, OFC-DH circuit(s) might mediate the contextual learning bias (or schema assimilation).

The ability of animals to form and use context-dependent, hierarchical associative structures (or context-informed cognitive maps) is arguably the foundation of high-level cognition^48,49^. This contextual knowledge contributes to the regulation of elicited behavior and influences learning. Using a bidirectional occasion-setting preparation, we show here that the orbitofrontal cortex is critical to all these manifestations of contextual knowledge. Moreover, our data suggest that different OFC circuits mediate the contextual regulation of elicited behavior and the contextual learning bias, the latter also involving the DH. Finally, this work highlights the importance of prior knowledge in learning, and provides a platform to investigate how new information makes contact with existing knowledge.

## METHODS

### Subjects

Male and female Long-Evans rats (Charles Rivers), 8-10 weeks at arrival, were housed in same-sex pairs in a temperature-controlled vivarium, with a 12-hr light/ dark cycle. Rats were mildly food-restricted to maintain∼95% of age-matched free-feeding weight, except during the pre-and post-surgery recovery period during which food was available ad libitum. Water was always available ad libitum in the home cage. Behavioral experiments were conducted during the light cycle. All experimental procedures were conducted in accordance with UCSB Institutional Animal Care and Use Committees and the US National Institute of Health guidelines.

### Apparatus

Behavioral training was conducted in 12 identical conditioning chambers, enclosed in individual sound-attenuating cubicles (Med Associates). A fan mounted on the cubicle provided ventilation and low background noise. Each chamber was equipped with four auditory cue devices; front panel: clicker and 4.5 kHz tone speaker; back panel: white noise and 2.9 kHz tone speaker (∼76dB each). Two ceiling-facing lights located on the front and back internal walls of the cubicle provided diffuse chamber illumination and were used to manipulate the background visual context. Sucrose solution (15% w/v) was delivered in a recessed reward port, located in center of the front panel, via a syringe pump located outside the sound-attenuating cubicle. Head entry into the reward port was detected by interruption of an infrared beam. Auditory cues were synchronized across boxes during behavioral sessions.

### Discrimination Training

Behavioral tasks were adapted from *Delamater et al*.^50^ Rats were first trained to consume sucrose from the reward port, in a single magazine training session (60 deliveries over 1h; 0.1ml/delivery). Rats were then trained daily (5-6 days/week) on one of two Pavlovian discrimination tasks: context-dependent (n = 48) or simple discrimination (n = 33). During the 80-min training sessions, the visual background (the context) alternated every two minutes between context A (flashing chamber lights, 0.25s on, 0.25s off) and context noA (all lights off). Within each context phase, two 10-s auditory cues, X or Y (white noise or 2.5Hz clicker, counterbalanced), were presented semi-randomly in two separate trials (80 trials per session; ITI = 50±20s), and potentially immediately followed by sucrose reward (0.1ml delivered over 3s). In the context-dependent discrimination task, the reward-predictive validity of each (auditory) cue was informed by the background (visual) context. Specifically, cue X was rewarded only in the presence of the context A, while cue Y was rewarded only in the absence of context A (A:X+ / X- / A:Y- / Y+). In the simple discrimination, one cue (X) was always rewarded whereas the other cue (Y) was never rewarded; the background context was irrelevant (A:X+ / X+ / A:Y- / Y-). The identity of the cue presented in a trial was selected semi-randomly so that the outcome of one trial did not inform the outcome of the next trial. During the first half of the pre-surgery training period (40 or 10 sessions for context-dependent or simple discrimination, respectively), rats experienced an equal number of rewarded and unrewarded trials per session. For the remainder of the experiment, the proportion of rewarded trials was reduced (20 rewarded vs. 60 unrewarded trials per session, unless specified otherwise) in an effort to promote discrimination accuracy, particularly in rats trained in the context-dependent discrimination. Rewarded trials were always equally distributed across contexts.

### Stereotaxic surgeries

Rats were anesthetized with isoflurane (induction: 4%; maintenance: 1–2.5%) and received either AAV8-hSyn-hM4Di-mCherry (Addgene #50475-AAV8) or AAV8-hSyn-mCherry (Addgene #114472-AAV8), bilaterally into the OFC or the DH. For the OFC, 0.5μL of virus was injected at the following coordinates: anterior/posterior (AP): 4.0, 4.0, 3.2, 3.2 from bregma; medial/lateral (ML): ±2.5, 2.5, 3.0, 3.0 from midline; dorsal/ventral (DV) -4.5, -4.4, -5.5, -5.4 from skull surface (coordinates for injection 1, 2, 3, 4, respectively). For the DH, 0.6μL of virus was injected at the following coordinates: ML:-3.4, -4.0, -4.6; ML: ±2.2, 2.6, 3.0; ML: -2.6, -3.0, -3.5 (coordinates for injection 1, 2, 3 respectively). Viruses were injected via 33 gauge needles (Hamilton #65460-03) connected to an infusion pump, at a rate of 1μL/min. Injecting needles remained in place for an additional 5min to allow for virus diffusion. The rats were allowed to recover for at least 1 week after stereotaxic surgery before resuming behavioral training. CNO-mediated chemogenetic silencing sessions were conducted >4 weeks post-surgery.

### Role of the OFC and DH in context-dependent reward-seeking (performance)

Following recovery from surgery, rats were returned to their respective discrimination training for an additional 16 sessions. During that period, rats were habituated to the injection procedure via daily mock i.p. injections (abdomen poked with a syringe, no needle attached). On separate days, rats then received i.p. injections of Clozapine-N-Oxide (CNO) dihydrochloride (HelloBio #HB6149, 5mg/kg, dissolved in saline) or saline (order of injection counterbalanced), 30min before being placed in the conditioning chamber for a standard behavioral session. Injections were separated by 2-3 regular sessions with no injections.

### Role of the OFC and DH in contextual learning bias

To assess the role of the OFC and DH in the knowledge-based inference of context-dependency during new learning, the same rats —previously trained in the context-dependent or simple discrimination task— were subjected to a new learning task. This phase of the experiment took place over 8 consecutive sessions (6 training sessions + 2 test sessions). Sessions 1, 3, and 5 consisted of the same context-dependent, or simple discrimination sessions, as previously described (each rat continued on the same task). Sessions 2, 4, and 6 were similar to these original training sessions, with the exception that a novel auditory cue (steady 4.5kHz tone overlaid with pulsating [0.5s on / 0.5s off] 2.9kHz tone, duration 10s) was introduced on some trials. For all rats, this novel cue (Z) was presented only in the context A, and always rewarded. These new (A:Z+) trials (10 per session) were interspersed with regular trials featuring the original cues and their regular programmed outcome (A:X / X / A:Y / Y trials). The proportion of the original trials was adjusted to maintain an equal number of reward deliveries in both contexts. This resulted in the following trial distribution: Ctx-dep: 5*[A:X+] + 45*[X-] + 45*[A:Y-] + 15*[Y+] +10*[A:Z+]; Simple discrimination: 5*[A:X+] + 15*[X+] + 45*[A:Y-] + 45*[Y-] + 10*[A:Z+]. Note the increased number of trials on these sessions (120 trials over 2h). All rats received CNO injections (5mg/ml, i.p.) 30min prior to the session introducing the new cue Z. Finally, the ability of the context to modulate responding to the different target cues was tested in two unrewarded probe tests (session 7 and 8). The first probe test (session 7, conducted 48h after the last CNO injection) assessed the contextual modulation of the new cue Z; the second probe test (session 8) assessed the contextual modulation of the original cues X and Y. During the probe tests, the target cues (X, Y, or Z) were presented three time in both contexts (test #1: A:Z / Z; test #2: A:X / X / A:Y / Y; trial order counterbalanced within each test session). No reward was delivered during the probe test sessions. Rats received no injection prior to the probe test sessions.

### Role of the OFC and DH in retrospective interference

Following test of inferred context-dependency, rats were returned to their original discrimination task (context-dependent or simple discrimination) for 5-6 sessions. Then, for the next 3 sessions only cue Z was presented, in both contexts, and always immediately followed by sucrose delivery (A:Z+ / Z+ trials; 20 trials per 80-min session). All rats received CNO injections (5mg/ml, i.p.) 30min prior to these sessions of consistent Z reinforcement. A final probe test (administered 48h after the last CNO injection) assessed discrimination accuracy on the original cues X and Y. This final probe test was a repeat of the previous probe test (A:X / X / A:Y / Y; 3 nonrewarded presentation of each cue/ context combination, 12 trials total). Data obtained on this final probe test (post-interference) were compared with data obtained on the previous probe test (pre-interference).

### Histological verification of DREADD expression

Rats were anesthetized with pentobarbital and transcardially perfused with PBS followed by 4% PFA. Brains were extracted, post-fixed in 4% PFA for 24h, cryoprotected in 30% sucrose for >3 days, and sectioned at 40μm on a freezing microtome. Coronal sections were collected onto glass slides and stained with a DAPI-containing mounting medium (Vectashield-DAPI, Vector Laboratories). Virus (mCherry) expression in the OFC or DH was examined under a fluorescence microscope (Keyence BZ-X800).

### Statistical Analysis

Cue-evoked reward seeking was quantified as the percentage of time a rat spent in the reward port during the last 5 seconds of cue presentation, when anticipatory magazine approach is more reliably observed and less contaminated by orienting behaviors51–53. To facilitate group comparisons, we calculated for each animal a discrimination index, defined as the difference between time-in-port during rewarded and nonrewarded trials divided by the sum. For the cue Z, we also calculated a contextual modulation index, defined as the difference between time-in-port during [A:Z] and [Z] divided by the sum. Statistical analyses were performed using SPSS statistical package (IBM SPSS, version 28). Specific analyses are described throughout the Results section, and compiled in Table S1. These analyses consisted generally of mixed-model ANOVAs with Cue, Context, and Injection as within-subject factors, and Task and Virus as between subject factors. Post-hoc comparisons were carried with Bonferroni-corrected t tests. Significance was assessed against a type I error rate of 0.05. Effect sizes were quantified by the partial eta squared (pη2) for ANOVAs main effects and interactions, and Cohen d for post-hoc comparisons. Six rats had to be excluded for failure to acquire context-dependent discrimination within 80 sessions; nine more rats had to be excluded for incorrect virus expression. The remaining rats were distributed as follows: Ctx-dep_OFC_hm4Di n=12 (7M + 5F); Ctx-dep_OFC_mCherry n=12 (7M + 5F); Simple_OFC_ hm4Di n=8 (5M + 3F); Simple_OFC_mCherry n=8 (4M + 4F); Ctx-dep_DH_hm4Di n=12 (6M + 6F); Ctx-dep_ DH_mCherry n=12 (6M + 6F); Simple_DH_hm4Di n=9 (5M + 4F); Simple_DH_mCherry n=8 (4M + 4F). One rat trained in the context-dependent discrimination and expressing hM4di in the OFC (Ctx-dep_OFC_hm4Di) had to be excluded before the contextual learning bias testing because of hardware malfunction (leading to n = 11 for this second experiment). No significant effects of sex were found, therefore data for males and females were collapsed. Note however that the study was not powered to investigate sex differences in these tasks.

## SUPPLEMENTAL MATERIAL

**Fig. S1:**
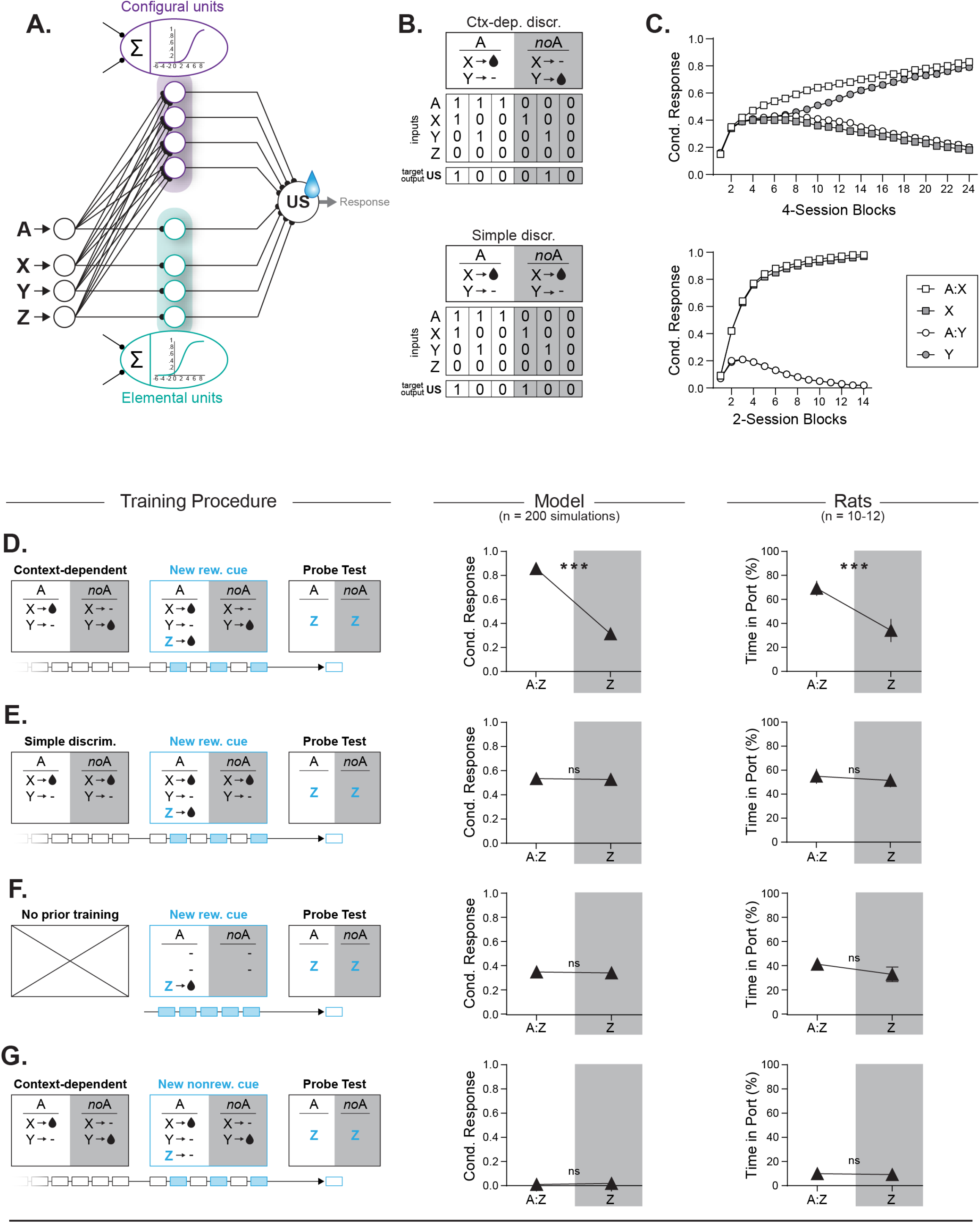
A simple connectionist model can account for context-gated reward predictions and the inference of context-dependency during new learning. **A**. Model description. The model (borrowing largely from [1] and [2]) consists of a simple feedforward neural network with three layers: one input layer (representing the different type of stimuli: contextual stimulus A and target stimuli X, Y or Z), one output layer (representing the reward expectation, or behavioral response), and one hidden layer connecting the inputs to the output. Units in the hidden layer are segregated in two categories: elemental units (receiving inputs from a single stimulus) or configural units (receiving converging inputs from all stimuli). Net activity in hidden units is converted to a propagated signal (ranging from 0 to 1) according to a logistic activation function. A constant non-adjustable negative bias, subtracted to the net activity of hidden units, ensures that these units remain relatively silent in absence of excitatory inputs. This negative bias is larger for the configural units (-5 vs. -2.5 in elemental units) resulting in a lower resting activity for these configural units (0.007 vs 0.07 for elemental units). As a result, the network initially favors elemental solutions during learning, and configural solutions are only engaged later in training if elemental approaches fail to accurately predict the reward (similar to what we observed in rats trained in our task). The weights of connections between units are initially randomized (between -1 and 1) and the classic backpropagation algorithm [3] was used to train the network (i.e. adjust the weights to minimize network error). Model parameters: learning rate: 0.08; momentum: 0.09. All simulations were conducted in Matlab R2020a. **B**. Example of behavioral tasks (context-dependent or simple discrimination) and corresponding inputs-targets outputs matrix used to train the networks. Note that the input for cue Z is always zero as this cue is not presented during initial training. **C**. Network performance during acquisition of context-dependent (top) or simple discrimination (bottom). On a given trial, the activation value for the US output unit corresponds to the reward prediction on this trial. To transform this reward prediction into an “observable behavioral response”, the US activation value (generally defined between 0 and 1 but occasionally slightly extending beyond this range) is clipped to the [0-1] range. Data show the average performance across 200 simulations. **D-G**. Effects of prior knowledge (i.e. prior discrimination training) on new associative learning in network simulations and animal subjects. *Left:* training histories. *Middle:* model simulations (n = 200). *Right:* empirical behavioral data (n = 11-12 surgery-naïve rats). Prior context-dependent discrimination training promotes the inference of context-dependency during new associative learning. Although the network initially favors learning via elemental units, prior training in context-dependent discrimination promotes the recruitment of configural units and allows for ctx.A (presented along the new cue Z during new learning) to “open the way” for Z to form an association with the reward US via the configural units. As a result, this newly formed association inherits context-dependency, as revealed in a final probe test in which responding to this cue Z is tested in both contexts (A:Z vs Z). This prediction was verified in animal subjects (**D**). This inferred context-dependency was not observed in networks or animals previously trained in simple discrimination (**E**), or in networks or animal that did not receive prior training (**F**). Moreover, networks or animals trained in the context-dependent discrimination and introduced to a new non-rewarded cue Z did not develop conditioned responding to that cue, regardless of context (**G**).*** P < 0.001. Error bars = s.e.m.

**Fig. S2:**
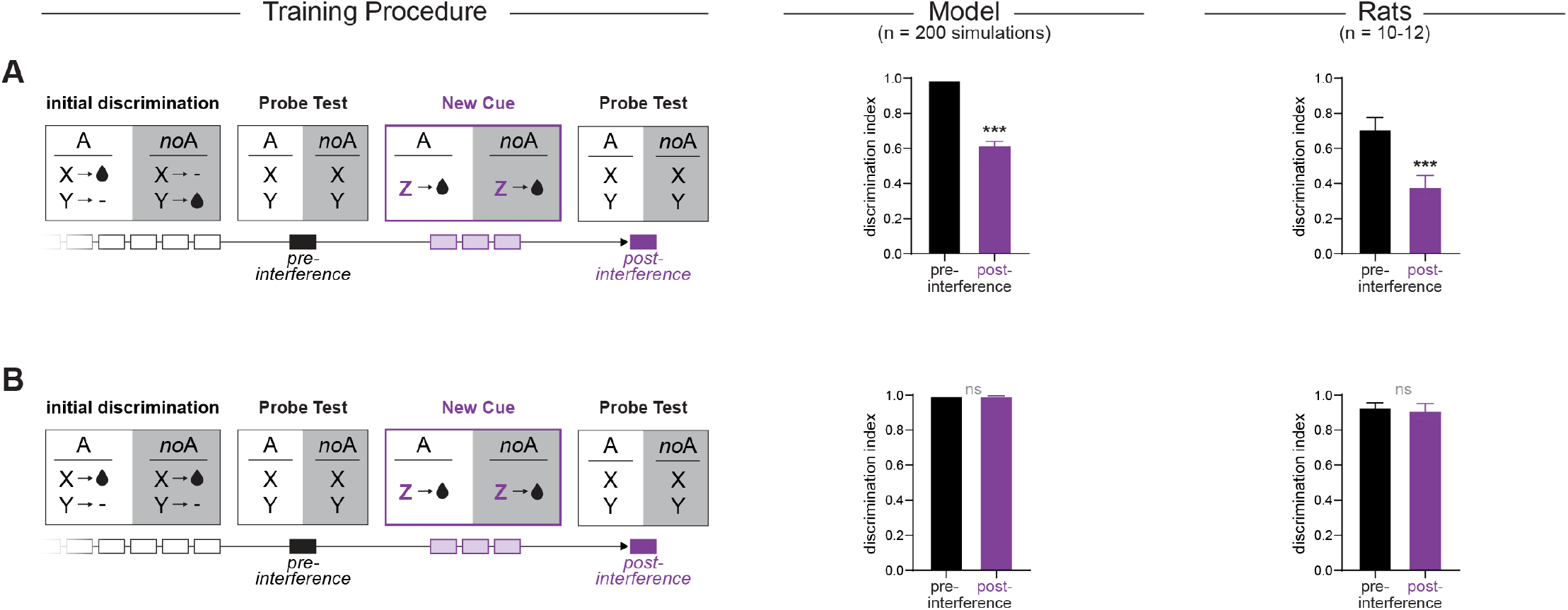
New context-independent associations decrease accuracy on original context-dependent discrimination (retroactive interference), in network simulations and animal subjects. *Left:* training histories. ANNs (same as previously described) or rats were trained in the context dependent discrimination (**A**) or the simple discrimination task (**B**). Following initial discrimination training, a new reward cue (Z) was introduced for 3 sessions. This new cue was always rewarded regardless of context (A:Z+ / Z+), contradicting the principle of context-dependency for rats trained in the context-dependent discrimination task. The effect of this new information on initial associative memories was evaluated in two probe tests (pre- and post-interference by Z). *Middle:* model simulations (n = 200). *Right:* empirical behavioral data (n =10-12 surgery-naïve rats). For ANNs/rats trained in the context-dependent discrimination, contradicting new information interferes with context-dependent associative memories, leading to a decrease in discrimination accuracy (retroactive interference). No disruption was observed for ANNs or rats previously trained in simple discrimination. ***: P<0.001 t -tests. Error bars = s.e.m.

**Fig. S3:**
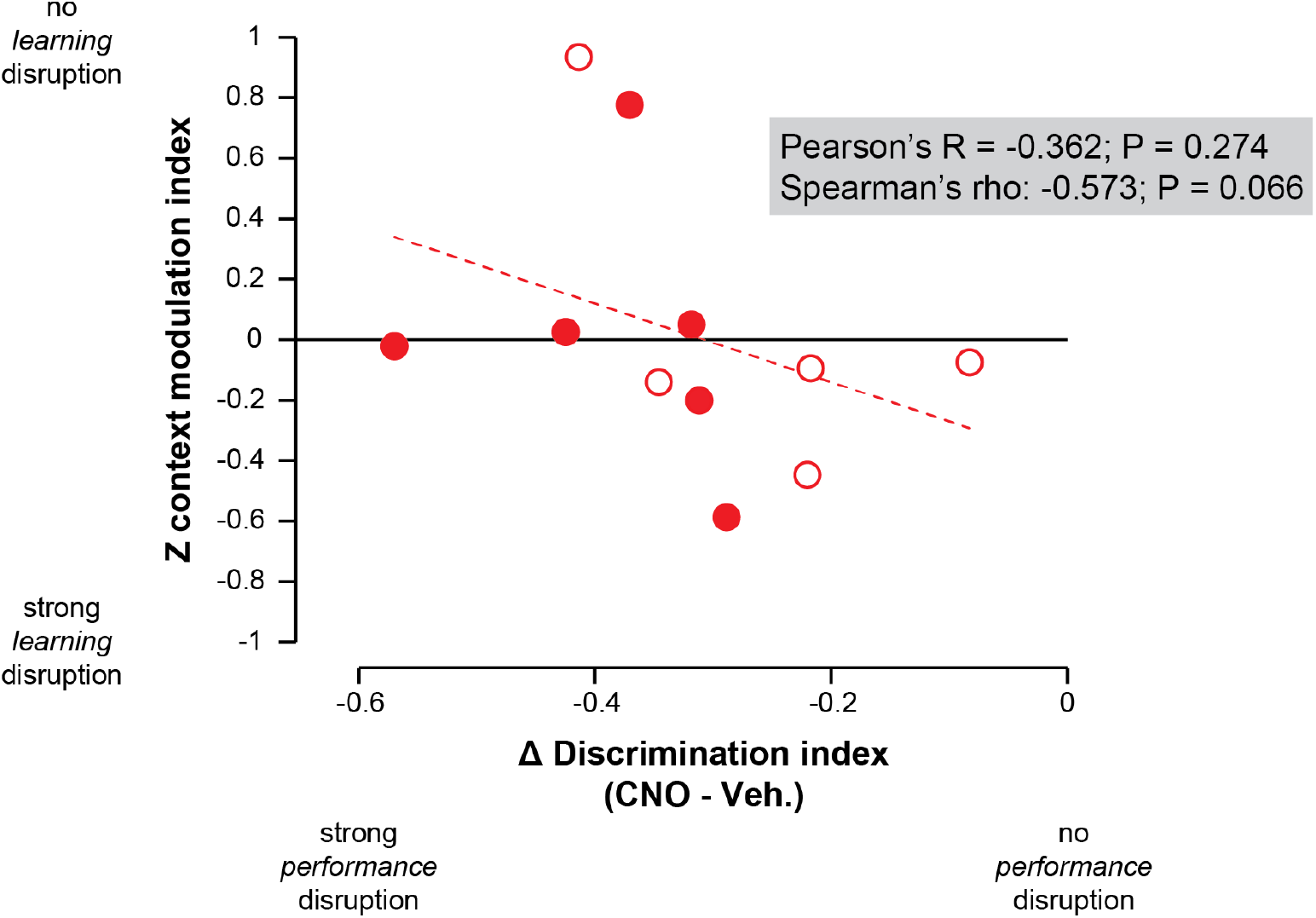
No correlation between the magnitude of OFC-dependent effects on context-dependent performance and contextual learning bias. Changes in the discrimination index between CNO and vehicle sessions (i.e., effect of OFC inactivation on performance; horizontal axis) and cue Z contextual modulation index (i.e., effect of OFC inactivation on contextual learning bias; vertical axis) for rats trained in a context-dependent discrimination task and expressing hM4Di in the OFC. Filled symbols: males; empty symbols: females.

**Table S1:**
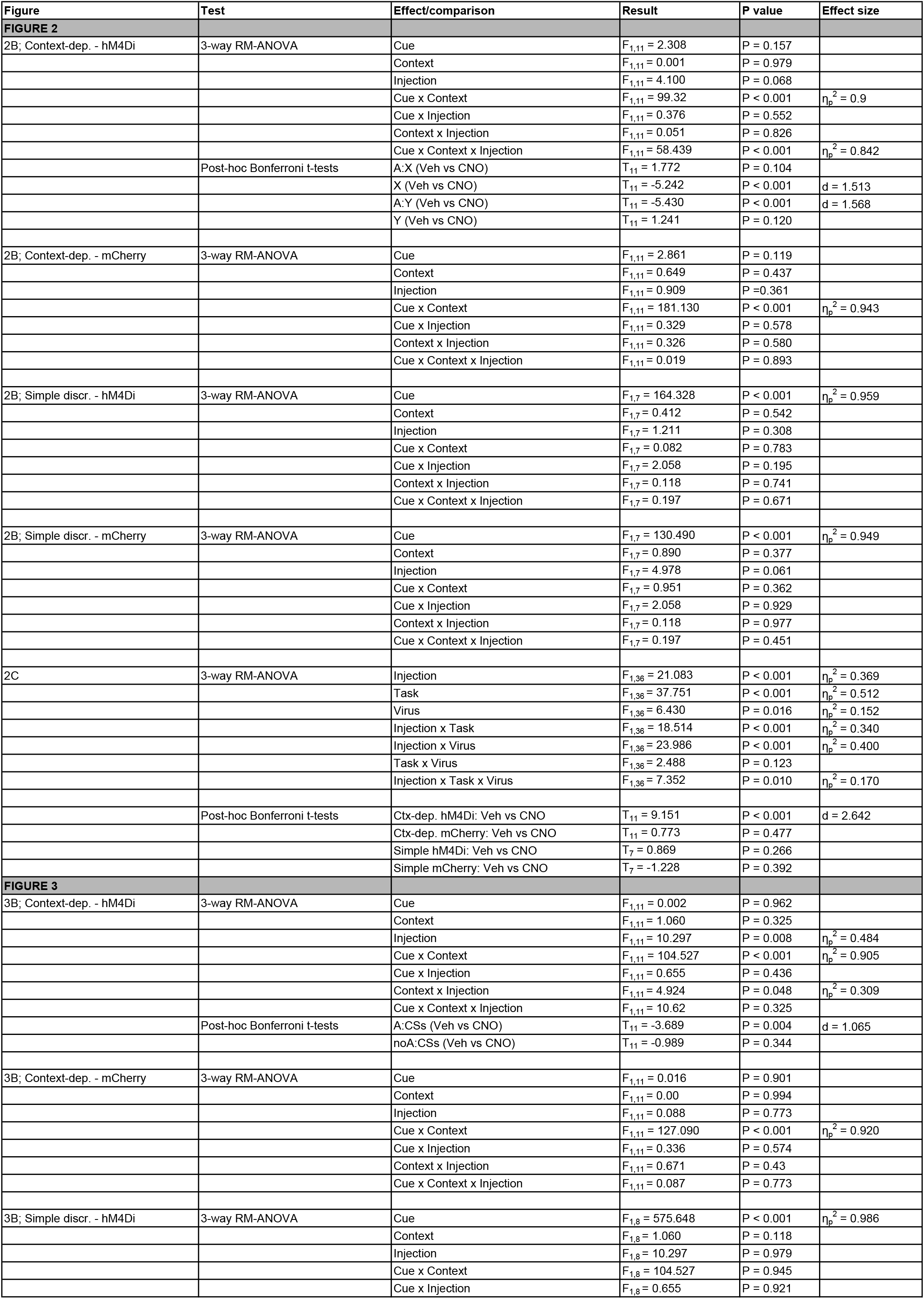
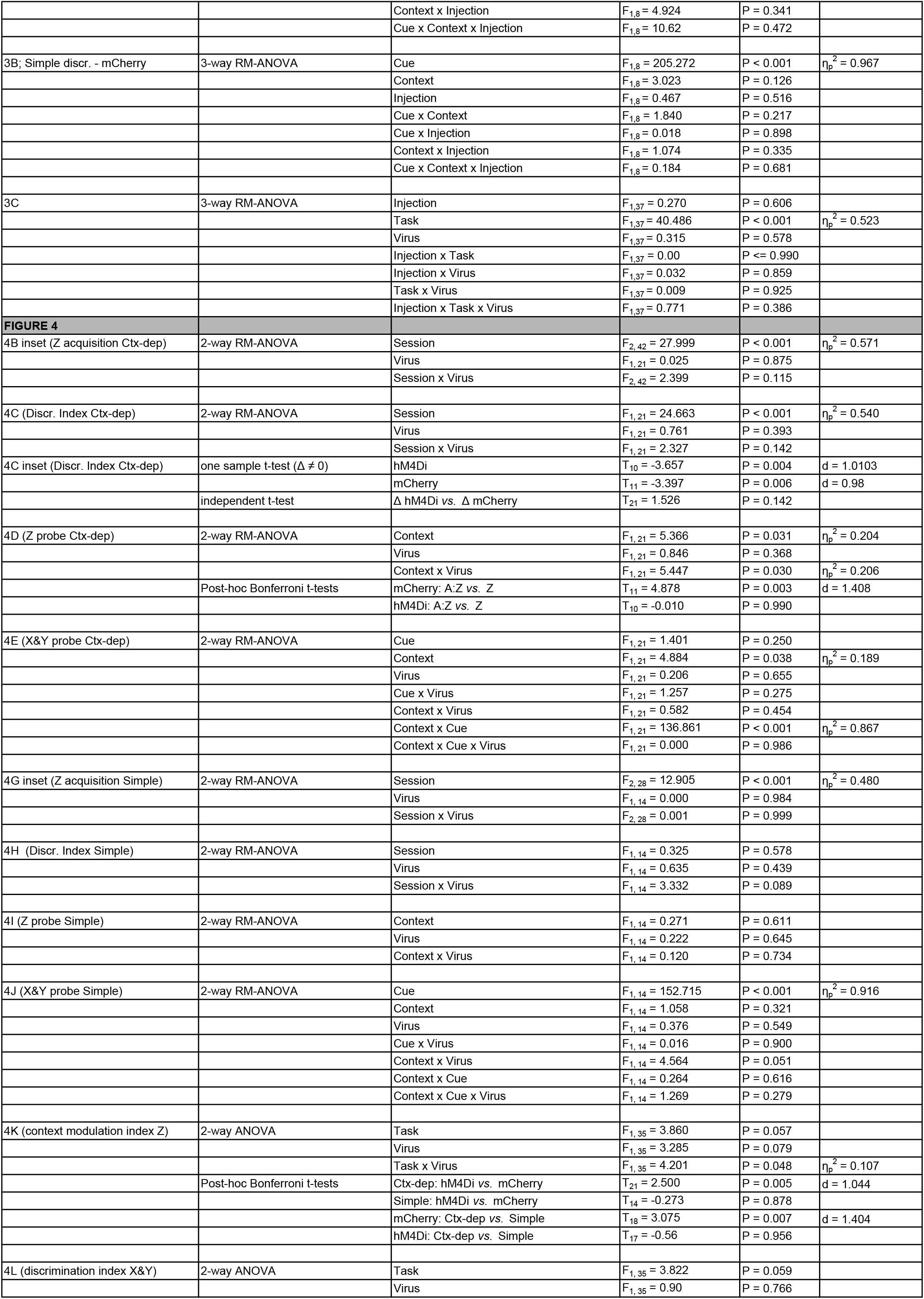
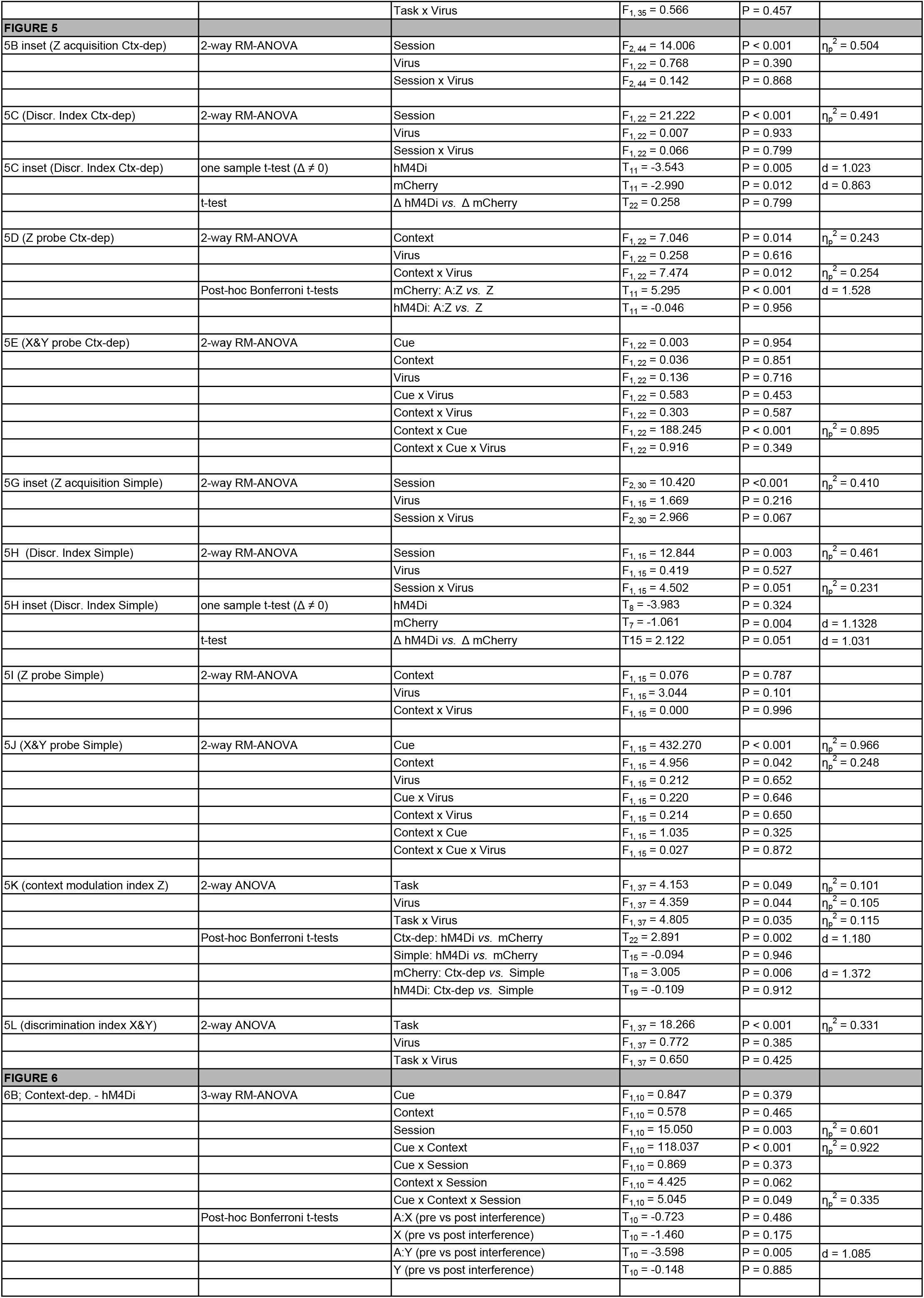
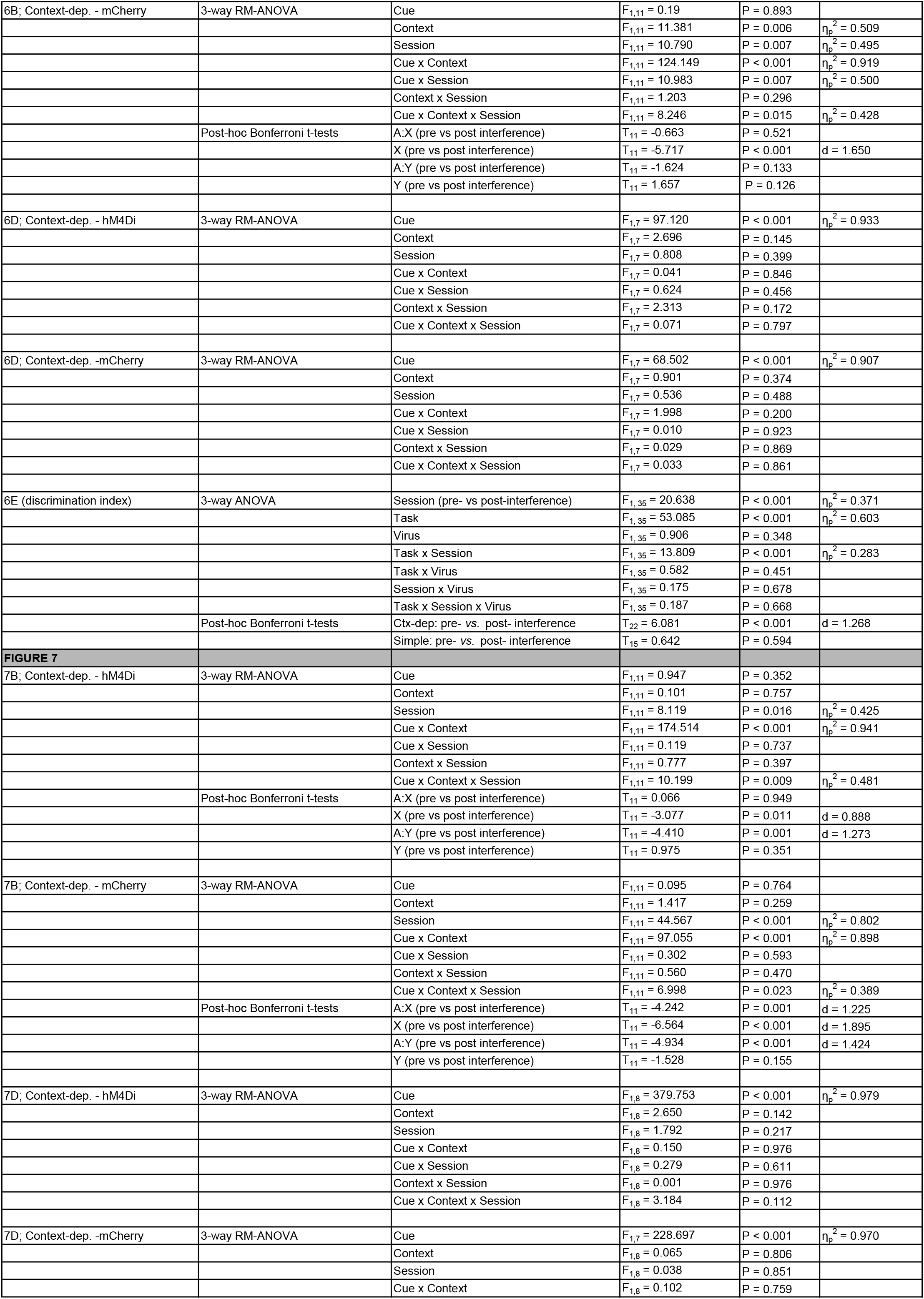
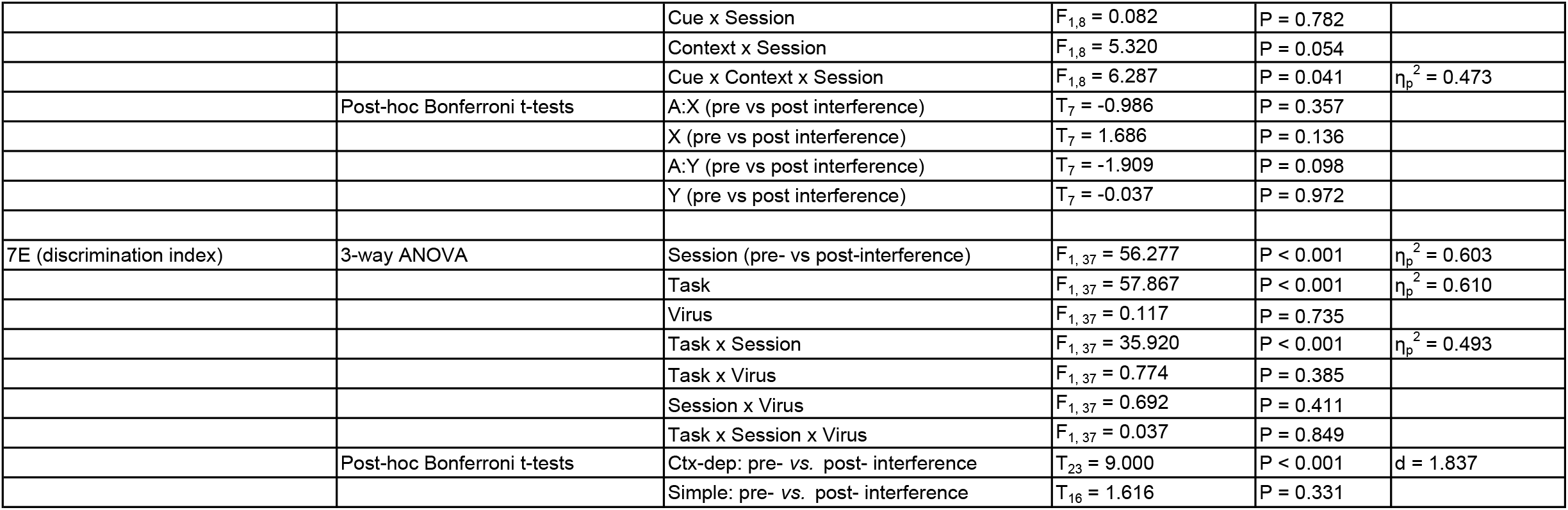
Comprehensive statistical results.

